# Associatively-mediated suppression of corticospinal excitability: A transcranial magnetic stimulation (TMS) study

**DOI:** 10.1101/600163

**Authors:** Manuel S. Seet, Evan J. Livesey, Justin A. Harris

**Author notes:** Address for correspondence: Manuel Seet, School of Psychology, University of Sydney, Sydney 2006, Australia.

## Abstract

Response inhibition—the suppression of prepotent behaviours when they are inappropriate— has been thought to rely on executive control. Against this received wisdom, it has been argued that external cues repeatedly associated with response inhibition can come to trigger response inhibition automatically without top-down command. The current project endeavoured to provide evidence for associatively-mediated motor inhibition. We tested the hypothesis that stop-associated stimuli can, in a bottom-up fashion, directly activate inhibitory mechanisms in the motor cortex. Human subjects were first trained on a stop-signal task. Once trained, the subjects received transcranial magnetic stimulation applied over their primary motor cortex during passive observation of either the stop signal (i.e. without any need to stop a response) or an equally familiar control stimulus never associated with stopping. Analysis of motor-evoked potentials showed that corticospinal excitability was reduced during exposure to the stop signal, which likely involved stimulus-driven activation of intracortical GABAergic interneurons. This result offers evidence for the argument that, through associative learning, stop-associated stimuli can engage local inhibitory processes at the level of the motor cortex.

## Introduction

Inhibitory control is considered a core mechanism underpinning our ability to manage behaviour and mental processes in service of attaining our goals. One popular paradigm used to assess inhibitory control is the stop-signal task (Lappin & Eriksen, 1966; Logan & Cowan, 1984). In a typical version of the task, the presentation of a go cue instructs subjects to perform a speeded motor response, such as a key-press. Occasionally, a stop signal is presented at a short delay after the onset of the go cue, in which case the initiated response has to be cancelled. The prevailing “horse-race” model construes stop-signal task performance as a function of two independent processes in competition with each other (Logan & Cowan, 1984; Verbruggen & Logan, 2009b). The go cue launches the go process to execute the key-press, and the stop signal triggers the stop process to suppress the ongoing response. If the go-process is completed earlier than the stop-process, the response is incorrectly produced (a failure of inhibition). Conversely, if the stop-process finishes prior to the go-process, the response is successfully withheld. Under this framework, the success rate of stopping reflects the efficiency of the inhibitory control process. Poor performance on the stop-signal task relates to deficits in restraining impulsive behaviours in everyday life (Logan, Schachar, & Tannock, 1997).

Countermanding a response is conventionally assumed to be effortful, goal-directed, and achievable with executive control (e.g. Aron, 2007; Logan & Cowan, 1984). Verbruggen and Logan (2008) challenged this predominant view, arguing that it is possible for response inhibition to be, at least in part, instigated by familiar environmental cues without cognitive mediation. In other words, stimuli can be conditioned, via repeated pairings with the act of stopping, to suppress motor output reflexively. Verbruggen and Logan (2008) introduced a paradigm to examine how the associative history of stimuli can impact future responses to those stimuli. In a go/no-go task, participants were trained to press a key when they encounter the names of living things but to make no response when they encounter the names of non-living things, for example. In the test phase, these stimulus-response mappings were retained for the control group, but were switched for the experimental group such that participants now responded to the names of non-living things and not to the names of living things. At test, go reaction times (RTs) were markedly slower in the experimental group, compared to RTs of the control group. The slower RTs in the experimental group were attributed to response interference created by an association between the previous no-go stimuli and response inhibition that was acquired during the training phase.

Using this modified go/no-go task, Chiu, Aron, and Verbruggen (2012) used transcranial magnetic stimulation (TMS) to investigate corticospinal excitability changes when participants were presented with stimuli previously associated with response inhibition. Single pulses of TMS were applied over the primary motor cortex (M1) of participants in the experimental group, during presentations of the go and no-go cues in the test phase. Motor-evoked potentials (MEPs) were measured using electromyography (EMG) of the task-performing hand. The peak-to-peak amplitude of the MEP indexes the excitability of the targeted motor cortical region and the descending corticospinal tract at the time of stimulation (Siebner & Rothwell, 2003). The authors hypothesised that stimuli frequently associated with response inhibition should suppress corticospinal excitability. Consistent with their predictions, MEP amplitudes during the test session were significantly smaller 100 ms after the onset of the cues that had previously served as no-go cues than after the cues that had previously served as go cues (Chiu et al., 2012). The result was taken as evidence that inhibition-associated stimuli quickly and automatically suppress the activity of the motor system.

There are two aspects of the experiment by Chiu et al. (2012) that limit the strength of their conclusions. First, the motoric impact of the inhibitory no-go cue (associated with withholding the response) was compared to that of the excitatory go cue (associated with executing the response). Therefore, the observed difference in corticospinal excitability between the two cue types could be partly or even wholly attributed to a residual excitatory effect of the former go cue rather than an inhibitory effect of the former no-go cue. To isolate an effect of associatively driven motor inhibition requires the motor effect of a stop-cue to be compared with that of a neutral cue matched for exposure time but not associated with going or stopping. The second limitation of Chiu et al’s experiment is that corticospinal excitability was probed as participants were preparing to make a response to a presented cue (in the condition where the previous no-go cues were now go cues). The motor system performs a series of preparatory processes during response selection and preparation (Bestmann & Duque, 2016; Poole, Mather, Livesey, Harris, & Harris, 2018), and these may affect MEP amplitudes at early time-points after cue onset. The extraneous influence of motor preparation can be avoided by examining how stop-related cues modulate corticospinal excitability when participants are merely observing the presented cues without needing to make or stop any response. These issues were addressed in the present research.

### Using TMS to study response inhibition

Corticospinal excitability can be studied using a single-pulse TMS protocol, in which MEPs are elicited by a suprathreshold test pulse administered over M1. TMS can also be used to probe inhibitory GABAergic interneurons in M1. In a paired-pulse protocol, a subthreshold pre-pulse is delivered 1-5 ms prior to the suprathreshold test pulse, causing the amplitude of the elicited MEP to be smaller than if the test pulse were administered alone (Kujirai et al., 1993). This phenomenon is known as short-interval intracortical inhibition (SICI). By comparing the amplitudes of single-pulse and paired-pulse MEPs, the magnitude of SICI can be quantified as the extent to which MEP amplitudes are reduced by the pre-pulse (Kujirai et al., 1993).

TMS has been used to profile motor cortical activity during the stopping of an initiated action. In the stop-signal task, on trials in which there is no command to stop (i.e. go trials), corticospinal motor neurons are engaged, which then send excitatory signals to the effector muscle via the corticospinal tract (van den Wildenberg et al., 2009). At the same time, GABAergic interneurons are inhibited, so as to facilitate the build-up of corticospinal excitability in preparation for response execution (Reynolds & Ashby, 1999). On stop trials, the onset of the stop signal will launch the stop-process, triggering the re-engagement of the GABAergic interneurons and subsequent inhibition of the corticospinal motor neurons (van den Wildenberg et al., 2009). Recent work in our laboratory has shown that the strength of GABAergic inhibition measured using SICI is strongly related to stopping performance in the stop signal task (Chowdhury, Livesey, Blaszczynski, & Harris, 2018). In light of the argument by Verbruggen and Logan (2008), we hypothesised that mere exposure to stop-associated stimuli can activate the GABAergic interneurons, even when there is no response to inhibit. As yet, there are no published studies that directly test this hypothesis.

### The present study

This study tested whether mere exposure to a stop-associated cue can suppress corticospinal excitability, and investigated whether GABAergic inhibitory circuits in the motor cortex are activated during this event. Participants first engaged in a version of the stop-signal task in a training phase. They continuously monitored a sequence of random letters on the computer screen, and had to respond as soon as they saw the letter X (the GO cue) but inhibit their response if a stop signal appeared shortly after the onset of the X. In the subsequent test phase, participants continued with the stop-signal task, which now contained additional trials during which TMS was administered. There were three types of TMS trials: GO-STOP trials, Stop-signal trials, and Control-stimulus trials. On GO-STOP trials, the stop signal appeared shortly after the go cue and participants had to withhold the cued response, as they did in the training stage. These were considered ‘active trials’ because there was a deliberate need to stop an ongoing action; such trials were included to maintain the need to stop responses occasionally, which is required for automatic inhibition to manifest (Chiu & Aron, 2014). On Stop-signal trials, the stop signal was presented after the onset of a random non-imperative letter. On Control-stimulus trials, a control stimulus (that was matched to the stop signal in style of presentation but not associated with responding or stopping) was presented after the onset of a random non-imperative letter. These latter two trial types were considered ‘passive trials’ because participants were merely exposed to stimuli without needing to stop any response. TMS was applied over the primary motor area, and MEPs were measured in the first digit interosseous (FDI) muscle of the index finger of the task-relevant (dominant) hand. In all TMS trials, stimulation was delivered 200 ms after the onset of the stop signal or the control stimulus. This time-point was chosen for TMS delivery because, by this post-onset time, inhibitory interneurons would already be activated in service of inhibiting responses in the stop-signal task (van den Wildenberg et al., 2009). This should increase the likelihood of observing changes in intracortical inhibition during exposure to the stop signal (if it is true that stop-associated stimuli can automatically activate inhibitory mechanisms). Stimulation on every alternate block used single-pulse TMS—in which a single suprathreshold test pulse was applied—while every other block used paired-pulse TMS—in which a subthreshold pre-pulse was presented 3 ms before the test pulse.

To measure changes in corticospinal excitability, we compared the amplitudes of MEPs elicited by single pulses of TMS (spMEPs) between the experimental conditions. We predicted that spMEPs would be smaller on Stop-signal trials than on Control-stimulus trials, consistent with our primary hypothesis that corticospinal excitability is suppressed by the stop signal. To investigate whether local inhibitory mechanisms within motor cortex contribute to the predicted change in corticospinal excitability, we tested for an interaction between the effect of the stop signal and the expression of SICI. As evidence of SICI, we expected that MEPs elicited by paired-pulse TMS (ppMEPs) would be smaller than spMEPs (Kujirai et al., 1993). However, this difference between spMEPs and ppMEPs should itself differ between Stop-signal and Control-stimulus trials if the local GABAergic mechanisms responsible for SICI are also implicated in the effect of the stop signal on corticospinal excitability. There are two ways in which this interaction could manifest. First, the combination of the stop signal and paired-pulse TMS may have a superadditive effect (see Abrahamyan, Clifford, Arabzadeh, & Harris, 2011; Du et al., 2015) if, for example, the sight of the stop signal increases the excitability of GABAergic neurons and augments the magnitude of SICI induced by the TMS pre-pulse. Alternatively, the combined effect of the stop signal and SICI might be subadditive, as has been observed for the combined inhibitory effects of SICI and long-interval cortical inhibition (LICI) produced by a pre-pulse of TMS delivered 100 ms before the test pulse (see Sanger, Garg, & Chen, 2001). Thus, a subadditive interaction between the stop signal and SICI might suggest that the stop signal affects corticospinal excitability by engaging GABAB mechanisms in the motor cortex that have been implicated in LICI (McDonnell, Orekhov, & Ziemann, 2006; Werhahn, Kunesch, Noachtar, Benecke, & Classen, 1999). Both of these interactions between the stop signal and SICI can be distinguished from what would be expected if the TMS pre-pulse and the stop signal act on independent inhibitory mechanisms. In this case, the effects of SICI and the stop signal should be additive (see Ethier, Brizzi, Darling, & Capaday, 2006; Townsend, Paninski, & Lemon, 2006), such that the magnitude of SICI on Stop-signal trials should be similar in size to that observed on Control-stimulus trials.

## Method

### Participants

Forty-two first-year psychology students at the University of Sydney took part in this experiment in exchange for course credit. Written informed consent was obtained and participants were screened using a questionnaire set by the National Institute of Neurological Disorders and Stroke. Only participants who declared that they did not have a history of relevant neurological, physiological or psychiatric conditions, and reported normal or corrected-to-normal visual acuity, without deficiency of colour vision, continued to the experiment. The Edinburgh Handedness Inventory (Revised) was used to determine handedness (Oldfield, 1971; Williams, 2010). Out of the sample of 42 participants, data from fifteen participants were excluded from analyses for reasons described in the Data Analysis section. The remaining 27 participants (mean age 19.5 years, 23 female, 25 right-handed) formed the final sample.

### Apparatus and Stimuli

Visual stimuli were displayed on a Dell LCD monitor, set to a spatial resolution of 1920 × 1080 and a refresh rate of 60 Hz. The procedure and the delivery of TMS were controlled using PsychoPy (Peirce, 2007), Version 1.82, run on a PC. A loudspeaker presented auditory signals. Key-press responses were performed on a standard ASCII keyboard.

All letters were presented in white in uppercase Arial, with a height of 3.0° and width of 2.9° of visual angle at a viewing distance of 60 cm. Two pairs of parallel bars were created, one served as the stop signal and the other as the control stimulus. The bars of each pair were separated by 6.7°, within which a letter was centrally presented; each bar had a thickness of 1.4° and a length of 9.6°. One pair of bars was horizontal and yellow, the other was vertical and cyan. These colours were approximately matched in luminance. All visual stimuli were displayed against a plain black background. Two auditory stimuli served as feedback signals, each lasting 0.5 s. The correct-response signal was a rapidly rising chime, while the error signal was a single buzz.

### TMS Procedures

Magnetic stimulation was administered using a pair of MagStim™ 200 stimulators yoked to a BiStim^2^ module (Whitland, UK). TMS outputs were conveyed via a 70mm-diameter figure-of-eight coil placed tangentially on the scalp, with the handle pointing backwards and deviating from the midsagittal axis by 45°. After completing the screening questionnaire, participants were seated in a chair facing the monitor. An adjustable forehead and chin rest comfortably supported the head at a viewing distance of 60 cm. The coil was placed over the primary motor cortex contralateral to the dominant hand. Participants wore a customised Lycra cap which marked out the optimal position for TMS of the motor cortex. Before subjects began the test phase of the experiment, which included TMS, their resting motor threshold was determined using the Rossini–Rothwell method (Rossini et al., 2015). Once the resting motor threshold had been determined (when MEPs ≥ 50μV, for 5 out of 10 times), an adjustable articulated arm (Manfrotto™, Italy) held the coil in position for the remainder of the experiment. The mean resting motor threshold was 54.3% of maximum stimulator output (SD = 10.0).

### EMG Recordings

Three self-adhesive Ag/Ag-Cl gel electrodes (10-mm diameter) were placed onto the opisthenar surface of the dominant hand, over the FDI muscle. Before electrodes were attached, the skin was lightly exfoliated with an abrasive pad and then cleaned with an alcohol swab. The positive and negative electrodes were placed over the distal and proximal sections of the FDI, respectively. The reference electrode was positioned over the ulnar styloid process. Surgical tape secured electrodes if necessary. EMG signals were recorded by a PowerLab 26T data acquisition unit (ADInstruments, New Zealand), and were amplified, sampled at 3 kHz and displayed using the data analysis software LabChart (ADInstruments, New Zealand). Before TMS thresholding and testing, EMG activity was inspected during resting state and flexion of the index finger, to ensure that recordings were not contaminated by extraneous noise. All data were digitised and saved for offline analysis.

### Experimental Procedure

The experiment ran for approximately 90 min per participant. After checking for the presence of a motor twitch, participants read an instruction sheet before beginning Phase 1.

#### Phase 1: GO-only training

Letters were presented sequentially one after another, each lasting 600 ms, in the centre of the black screen. No two consecutive letters were identical. GO trials comprised a GO cue, the letter X, appearing in the letter stream. While the X was still on the screen, participants had to press the spacebar as quickly as possible with the index finger of their dominant hand. A timely response was followed by the correct feedback signal, while failure to respond within the 600-ms presentation window was followed by the error signal. Participants completed two blocks of GO-only training, with each block lasting a minute and containing eight GO trials.

#### Phase 2: Stop-signal training

This phase featured a stop-signal task. Three types of trials were embedded in the continuous letter stream: GO trials, GO-STOP trials, and Control-stimulus trials (Figure 1). In GO trials, the letter X warranted a rapid spacebar press, as per the previous phase. Note that the 600-ms window to respond in GO trials served to prevent participants from strategically slowing their responses in order to succeed in the stop-signal task. In GO-STOP trials, the X cue was presented first, then followed, after a short delay, by the onset of one pair of bars flanking the X (the stop signal). On such trials, participants had to refrain from responding. The temporal asynchrony between the onset of the X cue and the stop signal is the stop-signal delay (SSD). Three SSDs—50 ms, 100 ms and 150 ms—were used, each employed for an equal number of STOP trials in randomly intermixed order. On these trials, participants received error feedback if they incorrectly pressed the spacebar. No auditory signal was delivered if they successfully withheld their response. In Control-stimulus trials, the other pair of bars (the control stimulus) appeared shortly after the onset of a non-imperative letter (i.e. any letter except X), at the same three delays used in GO-STOP trials. No response was required in these trials; participants were merely exposed to the control stimulus. The letter in Control-stimulus trials changed from trial to trial. The blue and yellow bars served as the stop signal and control stimulus in a counterbalanced fashion between participants. Participants completed 10 blocks of trials, with a cumulative running time of approximately 26 min, excluding breaks between blocks. Each block consisted of 15 GO trials, 6 GO-STOP trials, and 6 Control-stimulus trials. Trials were pseudo-randomised such that (i) the three trial types were distributed evenly throughout the block, and (ii) no more than three trials of the same type occurred successively. Consecutive trials were separated by either 5, 8 or 11 filler letters chosen at random.

**Figure 1.**
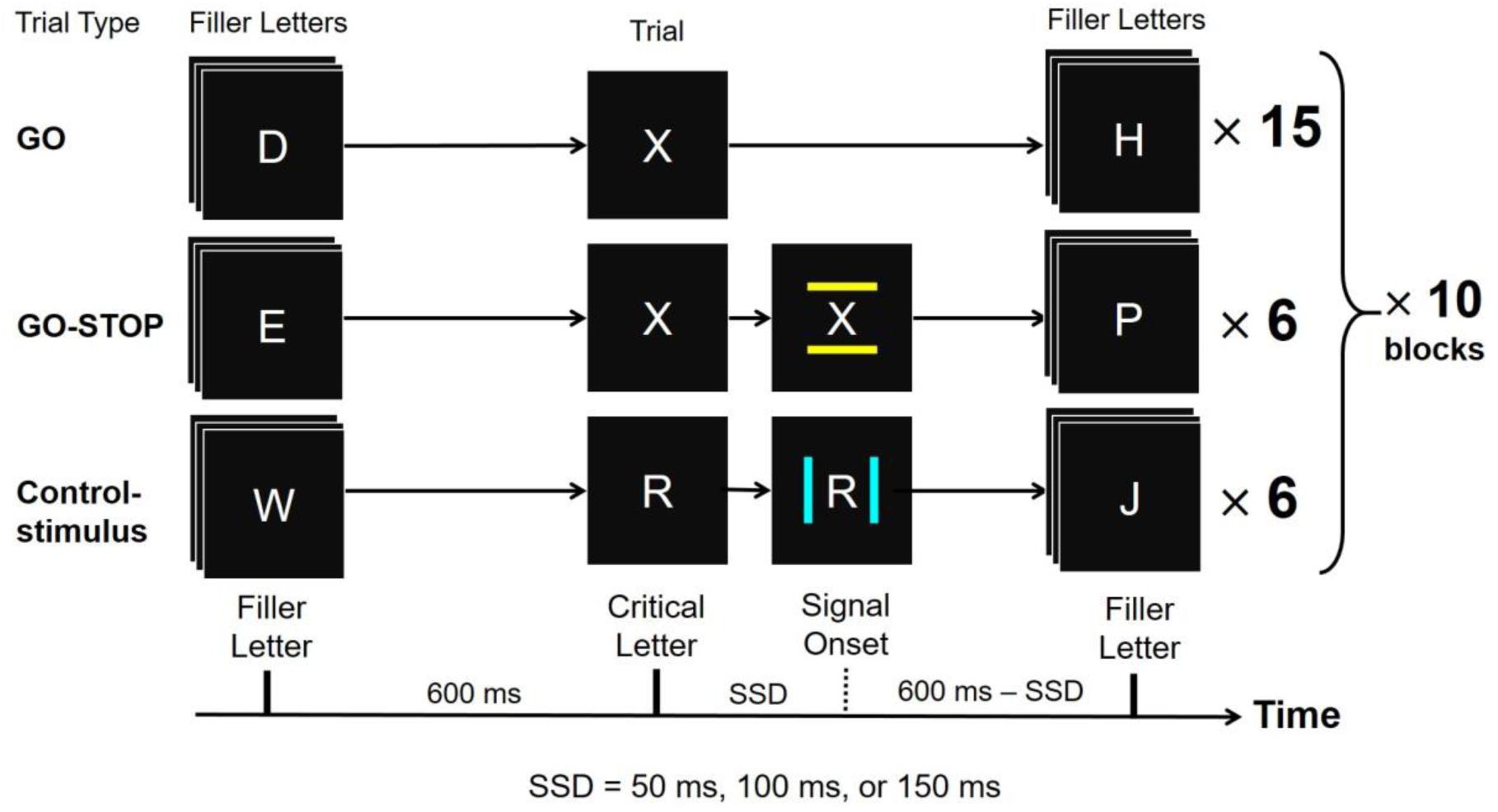
Design of the stop-signal training phase (Phase 2). In GO trials, the letter X instructed participants to press the spacebar. In GO-STOP trials, a stop-signal appeared shortly after onset of the letter X, instructing participants to withhold their response. Three SSDs were used with equal frequency in each block: 50 ms, 100 ms, and 150 ms. In Control trials, the control stimulus appeared shortly after the onset of a random non-imperative letter (note: this letter changes from trial to trial) at the three delays used in GO-STOP trials. In this example, the yellow horizontal lines serve as the Stop-signal and the vertical blue lines serve as the Control stimulus, but this assignment was counterbalanced across participants. The timeline at the bottom of the figure details the flow of events in the trials.

#### Phase 3: TMS test

Before Phase 3, EMG electrodes were placed on the subject’s dominant hand and the TMS thresholding procedure was performed. Participants then performed a stop-signal task similar to that in Phase 2. There were 8 blocks in this phase. Each block contained randomly mixed sequences of 46 trials, comprising 26 non-TMS trials, and 20 TMS trials (Figure 2). Non-TMS trials were identical in format to those in Phase 2; each block included 20 GO trials, 3 GO-STOP trials, and 3 Control-stimulus trials. For the latter two trial types, the signals always appeared 150 ms after the onset of the centrally presented letter (i.e. the SSD was 150 ms). There were three types of TMS trials: 5 GO-STOP trials, 5 Stop-signal trials, and 10 Control-stimulus trials in each block. In GO-STOP trials, the stop signal appeared shortly after the onset of the letter X, instructing participants to withhold their response. In Stop-signal trials, the stop-signal appeared in the presence of a random non-imperative letter. In Control-stimulus trials, the control stimulus appeared in the presence of a random non-imperative letter. For all TMS trials, the stop signal or the control stimulus always appeared 50 ms after the centrally presented letter, and TMS was always delivered 200 ms after the onset of the relevant stimulus.

**Figure 2.**
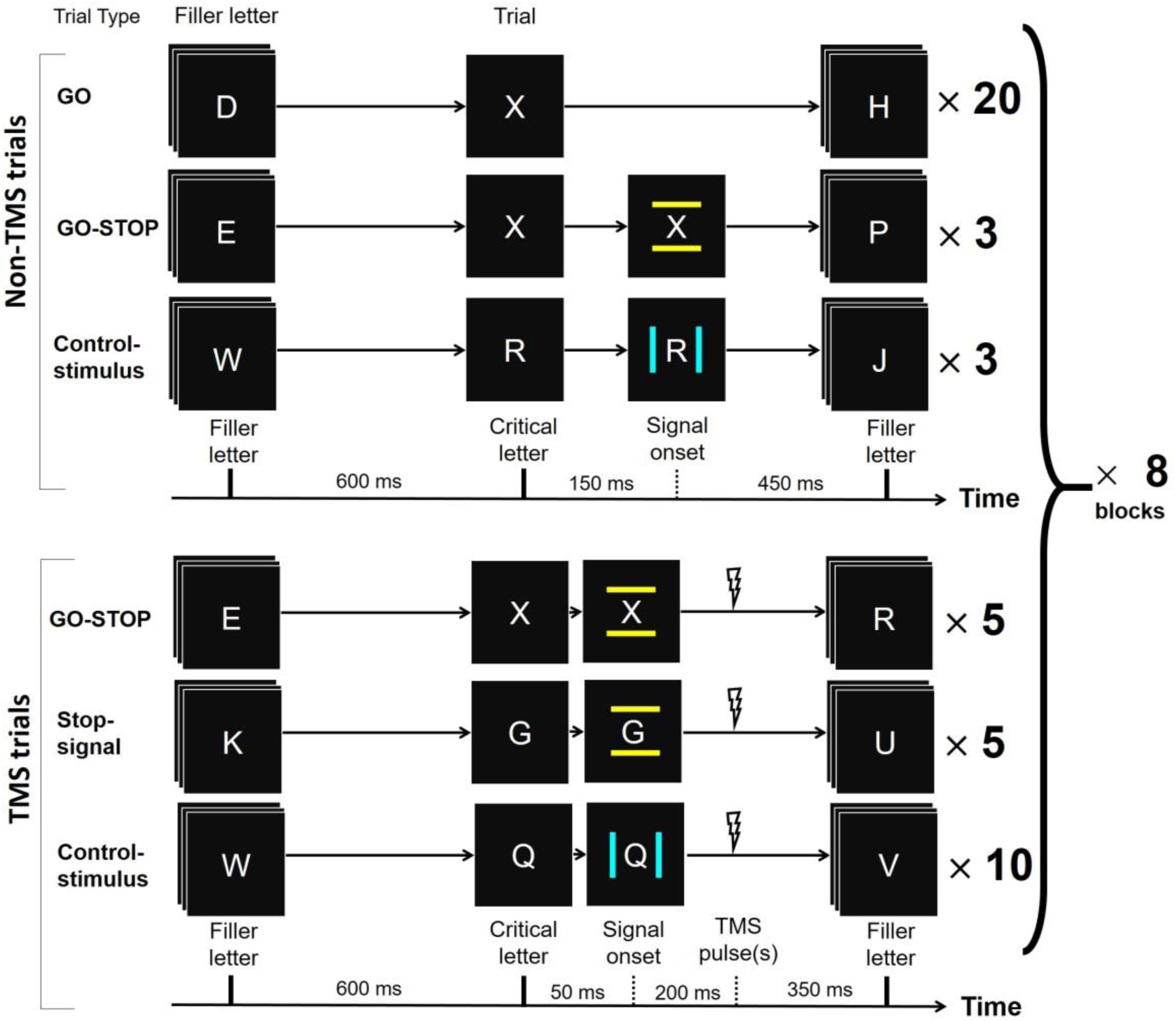
Design of the TMS test phase (Phase 3). The type of TMS delivered on TMS trials (i.e. single-pulse or paired-pulse) alternated between the blocks. In Stop-signal and Control-stimulus trials, the non-imperative letter contained between the bars changed from trial to trial. Note that all trials within each block were randomly intermixed.

Participants completed 8 blocks of trials, alternating between blocks using single-pulse TMS and those using paired-pulse TMS. In single-pulse TMS blocks, a single test pulse was delivered on TMS trials, set to an intensity which was 110% of the participant’s resting motor threshold. In paired-pulse TMS blocks, a pre-pulse—with an intensity which was 80% of resting motor threshold—preceded the 110% test pulse. The interval between the two pulses was 3 ms. The order of blocks (single-pulse vs paired-pulse) was counterbalanced across participants. That is, half of participants started with a single-pulse TMS block, while the other half began with a paired-pulse TMS block.

### Design and Data Analysis

MEPs were measured under three experimental conditions (GO-STOP, Stop-signal, and Control-stimulus trials). First, to test for changes in corticospinal excitability, we conducted a one-way ANOVA, followed up with a planned set of two orthogonal contrasts, on the spMEP amplitudes. To test our primary hypothesis that the stop signal reduces corticospinal excitability, the first planned contrast compared spMEPs between the Stop-signal trials and Control-stimulus trials. The second planned contrast compared MEP amplitudes between Active Response Trials (the GO-STOP trials) and Passive trials (averaging Stop-signal trials and Control-stimulus trials); this was done to test a secondary hypothesis about whether active motor preparation prior to successful inhibition can have an impact on corticospinal excitability. Second, to test for an interaction between SICI and the effect of the stop signal on MEPs, we conducted a two-way ANOVA comparing MEPs between two Pulse Number conditions (single-pulse vs paired-pulse) across three Trial Types (GO-STOP, Stop-signal, and Control-stimulus trials). This was followed by an ANOVA testing the same set of planned orthogonal contrasts on the 3-level factor of Trial Type and its interaction with the two-level factor of Pulse Number. Third, it is common in experiments on SICI to express the size of the inhibitory effect as the ratio of the ppMEP over the spMEP, such that a smaller value indicates a larger inhibitory effect (e.g. Roshan, Paradiso, & Chen, 2003). In keeping with this convention, we also conducted an analysis on these SICI ratios.

Out of the 42 subjects who participated in the experiment, data from five were discarded because they did not complete the test phase due to time constraints or technical problems. Data from one other participant were removed owing to excessive head movements, which prevented accurate targeting of TMS. MEPs of the remaining 36 participants were then screened on the following criteria. First, to ensure there was no significant pre-EMG activity that might contaminate with the upcoming MEP, the difference between the highest and lowest point of any waveform could not be larger than 100 μV within a 100 ms window preceding the TMS pulse; this criterion is identical to that adopted by Chiu et al. (2012). Second, we excluded data from failed stop trials to remove MEPs which were potentially affected by excessive muscle activity during response execution. Following these criteria, data from another nine participants were excluded because more than 50% of the TMS trials of at least one condition were deemed invalid.

The final dataset comprised data from 27 participants. Their data were then analysed in the following manner. For Phase 2, the 10 blocks of the stop-signal training were divided into 5 epochs each comprising two blocks. The successful stopping rate for each epoch was computed to be the percentage of GO-STOP trials in which no key-press response was produced. For the valid trials of Phase 3, the peak-to-peak amplitude of the MEP was measured as the voltage difference between the maximum and minimum point of a single waveform.

## Results

### Phase 1 and 2: GO and stop-signal training

The average go RT was 398.1 ms (SD = 26.0) in Phase 1. The go RTs and successful stopping rates over the 5 epochs in Phase 2 are plotted in Figure 3A, with within-subject standard errors of the mean (SEM) calculated according to the procedures of Loftus and Masson (1994). A one-way repeated-measures ANOVA revealed that the success rate of stopping was not equivalent across all epochs, 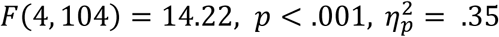. Orthogonal trend analyses showed that there was a significant linear trend, 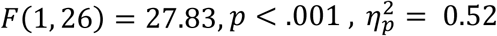, and a significant quadratic trend across epochs, *F*(1, 26) = 15.87, 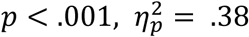. This confirms that stopping success improved over the course of the training.

**Figure 3.**
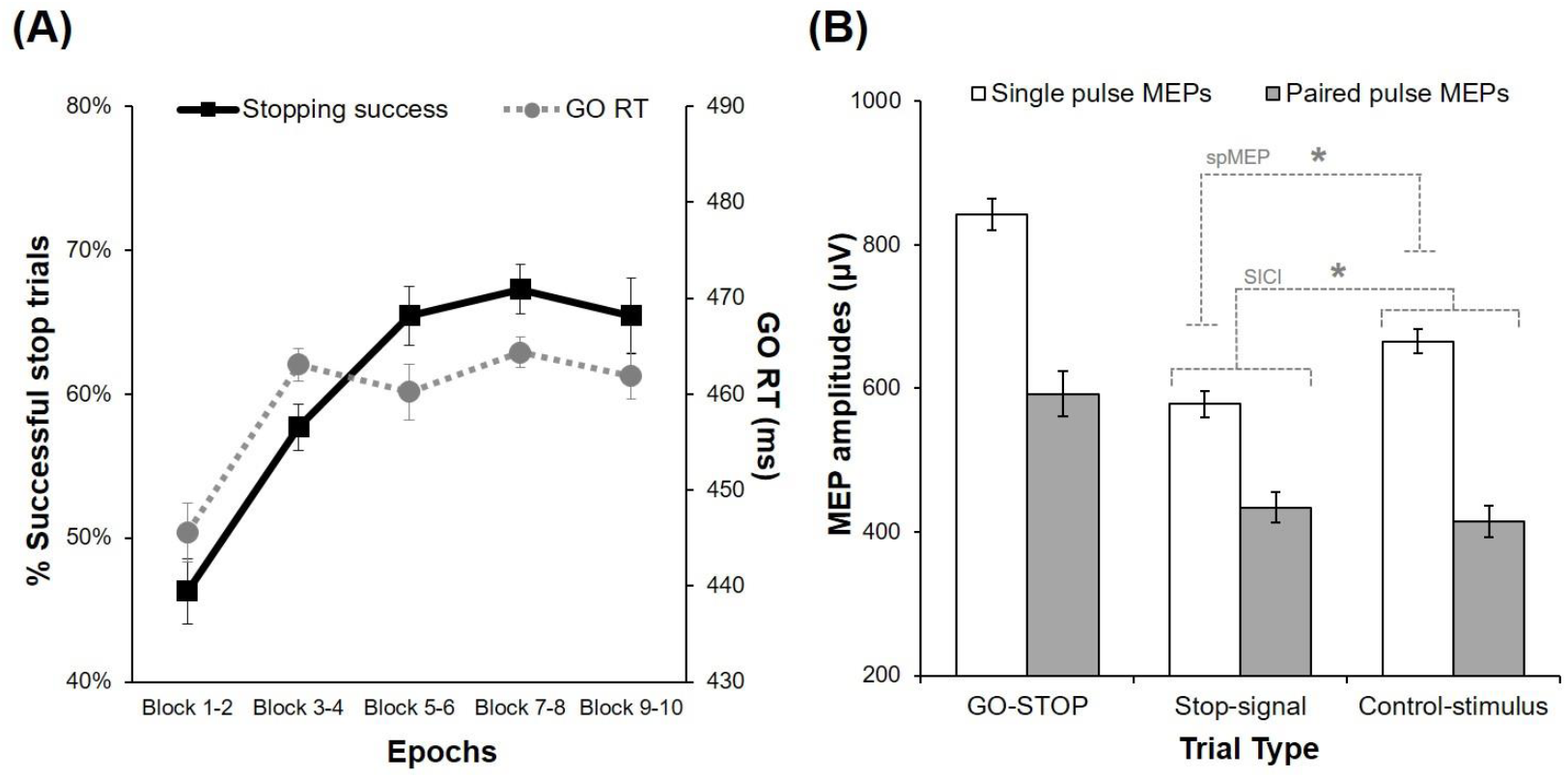
(A) Rate of successful stopping and GO RTs across the 5 two-block epochs of the stop-signal training phase (Phase 2). (B) MEP amplitudes in μV, as a function of Trial Type and Pulse Number, in the TMS test phase (Phase 3). Statistical significance of the difference in spMEP and SICI, between Stop-signal and Control-stimulus trials, are indicated (*p* <.05). All error bars represent ±1 within-subject SEM.

Improvements in stopping success can occur if participants slow their go RTs in anticipation of the stop signal (Verbruggen & Logan, 2009c). Indeed, the data shown in Figure 3A suggest that strategic slowing of go RTs may have contributed to the early improvement in stopping success observed between the first and second epoch. Nonetheless, strategic slowing cannot account for the continued improvements in stopping success over the remaining epochs. That is, while go RTs increased markedly from Epoch 1 to Epoch 2 (by an average of 17.5 ms), they remained stable thereafter, with no significant linear increase or decrease in go RTs across the 4 epochs from Epoch 2 through to Epoch 5, *F*(1, 26) < 0.01, *p* =.995. Meanwhile, stopping success rate continued to rise linearly during these same 4 epochs, 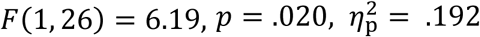. To investigate further whether strategic slowing could explain the rise in stopping success from Epochs 2-5, we calculated for each participant the linear slope of the change in their go RTs and the linear slope of the change in their stopping success rate across Epochs 2-5. The slope of stopping success was not significantly correlated with the slope of go RTs, *r*(25) = –.07, *p* =.742. This indicates that response slowing does not account for the observed improvements in stopping success. Thus improvements in inhibition across training appear to be a consequence of initial proactive control adjustments (strategic slowing) early in training in combination with continued improvements as a consequence of experience with the task and stimuli. The latter of these would be consistent with the notion of inhibition becoming automatically elicited by the stop signal.

### Phase 3: TMS Test

The average go RT was 475.1 ms (SD = 18.6) in Phase 3. The mean MEP amplitudes for each trial type were calculated, and the average of these across participants is depicted in Figure 3B. First, a one-way ANOVA was performed to compare spMEP amplitudes across the three trial types, and Greenhouse-Geisser corrections were used for cases in which the sphericity assumption was violated. There was a significant main effect of Trial Type,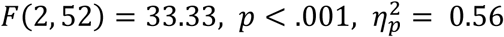. In the subsequent set of planned orthogonal contrasts (with α =.05), the first contrast revealed that spMEPs were smaller on Stop-signal trials than on Control-stimulus trials,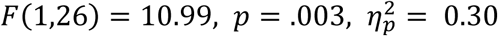. The second contrast found that spMEPs were larger on Active Response Trials than on Passive trials (Stop-signal and Control-stimulus trials),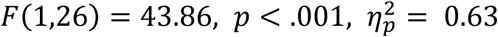.

Second, a (2× 3) ANOVA was conducted to examine MEPs between Pulse Number across Trial Types. There was a significant main effect of Pulse Number, 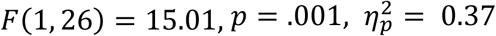. That is, averaging across all three trial types, MEP amplitudes were smaller for ppMEPs than for spMEPs; this is evidence of SICI (Kujirai et al., 1993). The main effect of Trial type was significant,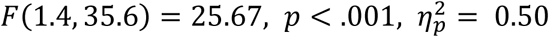, indicating that MEP amplitudes (averaging over Pulse Number) were not equal across the three types of trials. The Pulse Number × Trial Type interaction was also statistically significant, 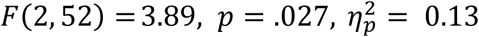. Next, planned orthogonal contrasts were used to compare MEPs between Stop-signal trials and Control stimulus trials, and between these “passive” trials with active (GO-STOP) trials. When averaged across Pulse Number, MEP amplitudes were not significantly different between Stop-signal trials and Control-stimulus trials,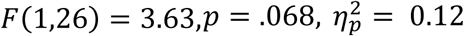, however the difference between these trial types did interact significantly with pulse number,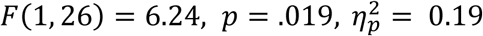. This interaction indicates that the difference between single- and paired-pulse MEPs was greater on Control-stimulus trials than on Stop-signal trials. Averaged over pulse number, MEP amplitudes were larger in Active trials than Passive trials (averaging Stop-signal trials and Control-stimulus trials),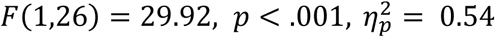, but this difference did not interact significantly with pulse number, 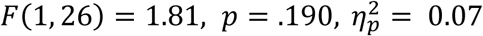.

Third, a one-way ANOVA examined SICI ratio data (ppMEP over spMEP) across the three Trial Types. It revealed that SICI was significantly different across Trial Types,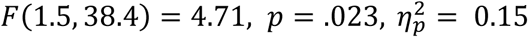. The two orthogonal contrasts tested previously showed that SICI was weaker for Stop-signal trials (mean ratio = 0.89) than for Control-signal trials 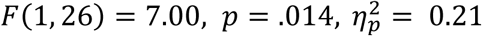, but SICI was not significantly different between Active trials (mean ratio = 0.71) and Passive trials 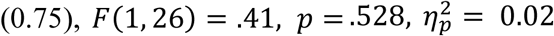. These results are consistent with the previous analyses performed on the differences between single- and paired-pulse MEPs.

## Discussion

The present experiment tested the proposition that stop-associated stimuli can automatically activate the same inhibitory mechanisms that are voluntarily used to interrupt a motor response. Averaging across single- and paired-pulse trials, amplitudes of MEPs were larger in active trials than in passive trials. Indeed, intentionally commanded processes (e.g. the go process) can significantly modulate corticospinal excitability even when no response is produced, and such motor preparatory modulations have been shown to be highly variable from trial to trial (e.g. Klein-Flügge, Nobbs, Pitcher, & Bestmann, 2013). This highlights why it was not optimal that Chiu et al. (2012) assessed stimulus-driven motor effects during trials in which a potential response had to be made or prevented.

To partial out any extraneous influence of voluntary motor processes, it was best to examine MEPs when participants passively observed the stop signal without needing to stop any response. Mere exposure to the stop signal brought about a suppression in corticospinal excitability detected at 200 ms after signal onset, as revealed by single-pulse MEPs. This is similar to the reduction of corticospinal excitability which reportedly occurred during encounters of go-cues previously associated with withholding a response (Chiu et al., 2012). However, the present experiment has two theoretically important differences over past work. First, the stop signal was compared to a control stimulus matched for familiarity and manner of presentation in the task but not associated with going or stopping. This allowed us to isolate the conditioned motor effects of the stop-associated stimulus, with respect to an appropriate control condition. Second, this experiment looked at the motor effects of the stop signal during moments when no intentional preparatory motor processes were in progress. Such a design minimised significant modulations of motor cortical activity which would directly affect MEP amplitudes and contaminate evidence for stimulus-driven motor inhibition. This experiment demonstrated that passive observation of a stop-associated stimulus was sufficient to inhibit the motor cortex and corticospinal tract even when there was no need to stop any response at that time. This finding supports the ideas expressed by Verbruggen and Logan (2008). It must be noted that such automatic inhibition has been shown to manifest only in a context in which there is an occasional requirement to actively inhibit responses on other trials, in what has been referred to as an ‘executive setting’ (Chiu & Aron, 2014). Indeed, this executive setting was incorporated into the TMS test phase of the current experiment, by including GO-STOP trials with a relatively long 150-ms SSD (in which the average success stop rate is 50.0%, SD = 16.6%). Thus it is possible that the corticospinal suppression observed here would not be observed if no GO-STOP trials were present in the test phase. Given that this issue concerning the role of an executive setting in the manifestation of automatic inhibition has already been demonstrated in past research (Chiu & Aron, 2014; Verbruggen & Logan, 2009a), it is not of primary interest for the same issue to be tested in the current study.

The extent of SICI was smaller in the Stop-signal condition compared to the Control-stimulus condition. Such a pattern is inconsistent with there being independent inhibitory processes responsible for SICI and for the suppression of MEPs by the stop signal, since that would predict linear summation of these independent effects (see Ethier et al., 2006; Townsend et al., 2006) and thus equivalent size of SICI in both the Stop-signal and Control-stimulus conditions. The fact that the stop signal reduced the size of SICI suggests that the stop signal might affect corticospinal excitability by engaging the GABAB inhibitory mechanisms. This conclusion is based on evidence that SICI, which depends on GABAA receptors (Di Lazzaro et al., 2000), is itself suppressed when LICI, which depends on GABAB receptors (McDonnell et al., 2006; Werhahn et al., 1999), is engaged (Sanger et al., 2001). However, the fact that SICI was reduced in the presence of the stop signal could also mean that the stop signal and SICI act via the same GABAA receptors to reduce corticospinal excitability. In this case, the subadditive interaction could occur if the stop signal engages the inhibitory processes and thereby limits the opportunity for the TMS pre-pulse to drive the same mechanisms to produce SICI. This type of interaction is similar to the observation that SICI, induced by a pre-pulse, is not significantly increased if the GABAA interneurons were recently activated (Ziemann, Rothwell, & Ridding, 1996). It is plausible that both of these proposed accounts hold true in explaining the observed reduction in SICI. Regardless of the type of GABAergic mechanism involved, the stimulus-driven engagement of the inhibitory motor circuits appears to be occurring within the first 200 ms after the onset of the stop signal, thereby producing the suppression of single-pulse MEP amplitudes.

In conclusion, the current study offers evidence suggesting that the mere sight of visual cues associated with the act of stopping can automatically decrease the excitability of the corticospinal pathway. The fact that the effect was observed when the stop signal was presented without a GO cue means that the stop signal affected motor cortical excitability without the top-down influence of any decision to cancel an action. This suppressive effect is likely attributable to the stimulus-driven activation of GABAergic interneurons in M1—circuitries which are normally implicated in voluntary restraint of motor responses (van den Wildenberg et al., 2009). Such findings not only strengthen support for the notion of associatively-mediated inhibition proposed by Verbruggen and Logan (2008), but also bolster the general idea that perceptual information can be quickly and automatically transformed into their affiliated motor codes.

## Acknowledgements

This research was supported by funding from the Australian Research Council, Grant DP160102871

## Conflict of Interest

The authors declare that they have no conflict of interest.

